## Appendix for "N-ACT: An Interpretable Deep Learning Model for Automatic Cell Type and Salient Gene Identification"

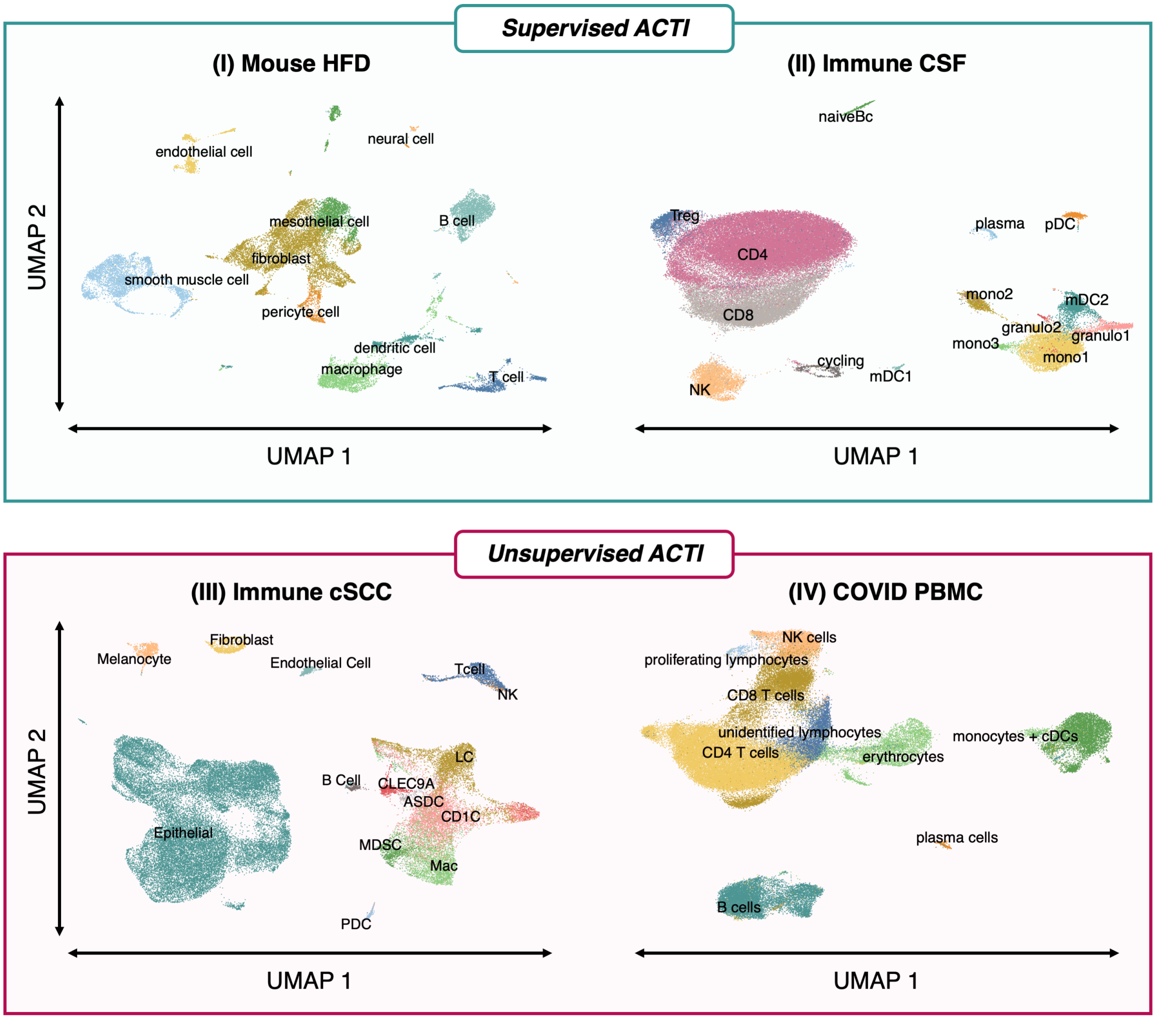

Figure A.3: A UMAP-reduced plot of cell populations present in each dataset investigated in this work.

#### B. 2.1. Mouse-HDF

Mouse-HDF (Kan et al., 2021) [SCP1361] consists of scRNAseq of aortic cells in mice given a normal diet and mice given a high-fat diet (HFD), resulting in 24K cells (24K cells after processing). The authors identified 27 clusters, for 10 different cell populations.

$$H(\mathbf{y}, \hat{\mathbf{y}}) = \sum_{j=1}^{k} y_j \log(\hat{y}_j) + (1 - y_j) \log(1 - \hat{y}_j), \tag{A.2}$$

where $\mathbf{y}$ and $\hat{\mathbf{y}}$ denote the original/generated labels and model predicted labels, respectively. In this formulation, $\mathbf{y}$ is a one-hot encoded vector of the cell types; that is

#### C. 5. F1 Score

F1 score is a standard metric for evaluating a classifier. F1 score is the harmonic mean of precision and recall, which is shown in Eq. (A.5),

$$F1 = 2\left(\frac{\text{precision} \cdot \text{recall}}{\text{precision} + \text{recall}}\right). \tag{A.5}$$

For more information on weighted and non-weighted F1 score, see `https://scikit-learn.org/stable/modules/generated/sklearn.metrics.f1_score.html`.

### Appendix D. CellMeSH Cell Type Predictions

CellMeSH (Mao et al., 2021) is a probabilistic method that generates a cell type prediction based on a gene list. CellMeSH aims to alleviate two common issues with querying gene lists from the literature: 1) publication bias, which relates to specific genes/cell types that are studied more often than other types, and therefore have more literature associated with them; and 2) Noise present in gene/cell-type mapping and the corresponding database. CellMesh uses a database built from the National Library of Medicine (NLM) MEDLINE indexed records called Medical Subject Headings. In building the CellMeSH database, Mao et al. use TF-IDF to reduce publication bias thus addressing publication bias.

To address the second issue, CellMeSH utilizes a probabilistic querying method. Given gene query list $Q$, CellMeSH assumes a probabilistic model for a query genes $g$ being obtained from a cell-type $C$, shown in Eq. (A.6):

$$P(g|C) = \begin{cases} \alpha \cdot w_C(g) & g \in Q \cap C \\ (1-\alpha) \cdot \frac{1}{N_g - K_C} & g \in Q \cap \overline{C} \end{cases} \tag{A.6}$$

with $w_C(g)$ being the adjusted weight of gene $g$ in cell type $C$ (using TF-IDF), $N_g, K_C$ denoting the total number of genes and total number genes with non-zero weight in $C$. $\alpha$ is a parameter which aims to help with noise in the database. For each candidate cell type $C$, CellMeSH calculates a log-likelihood score:

$$L(Q|C) = \log P(Q|C) = \sum_g \log P(g|C), \tag{A.7}$$

which are then used to rank $C$ based on their values. Lastly, CellMeSH uses a maximum likelihood-based estimation to generate predictions. In this framework, the top cell type, $C^*$, can be found as:

$$C^* = \text{argmax}_C \log L(Q|C). \tag{A.8}$$

### Appendix E. : Main Manuscript Results (Larger Figures)

**COVID PBMC**

| Population Annotation | CellMesh Hit | Top Retrieval |
|---|---|---|
| Natural Killer Cells | #1 | Natural Killer Cells |
| CD4+ T Cells | #5 | T Cells |
| Unidentified Lymphocytes | N/A | T Cells |
| B Cells | #1 | B Cells |
| Proliferating Lymphocytes | N/A | CD8+ T Cells |
| Monocytes + Dendritic Cells | #1 | Monocytes |
| Plasma Cells | #2 | B Cells |
| CD8+ T Cells | #2 | T Cells |
| Erythrocytes | #1 | Erythrocytes |

**Hit@1**: 0.57 **Hit@3**: 0.85 **Hit@5**: 1.00 **Hit@10**: 1.00

**Immune cSCC**

| Population Annotation | CellMesh Hit | Top Retrieval |
|---|---|---|
| Endothelial Cells | #1 | Endothelial Cells |
| Epithelial Cells | #2 | Keratinocytes |
| Fibroblast | #2 | Endothelial Cells |
| Melanocytes | #1 | Melanocytes |
| Plasmacytoid Dendritic Cells | #2 | T Cells |
| CLEC9A+ Dendritic Cells | #1 | Dendritic Cells |
| CD1c+ Dendritic Cells | #1 | Dendritic Cells |
| Macrophages | #1 | Macrophages |
| A+S+ Dendritic Cells | #1 | Dendritic Cells |
| T Cells | #1 | T Cells |
| MDS Cells | NR | Hematopoietic Stem Cells |
| B-Cells | #1 | B Cells |
| Langerhans Cells | NR | Reed-Sternberg Cells |
| Natural Killer Cells | #4 | T Cells |

**Hit@1**: 0.57 **Hit@3**: 0.78 **Hit@5**: 0.85 **Hit@10**: 0.85

Figure A.4: (For the ease of the reader) An enlarged version of Hit@K results, initially presented in Fig. 2 of the main manuscript.

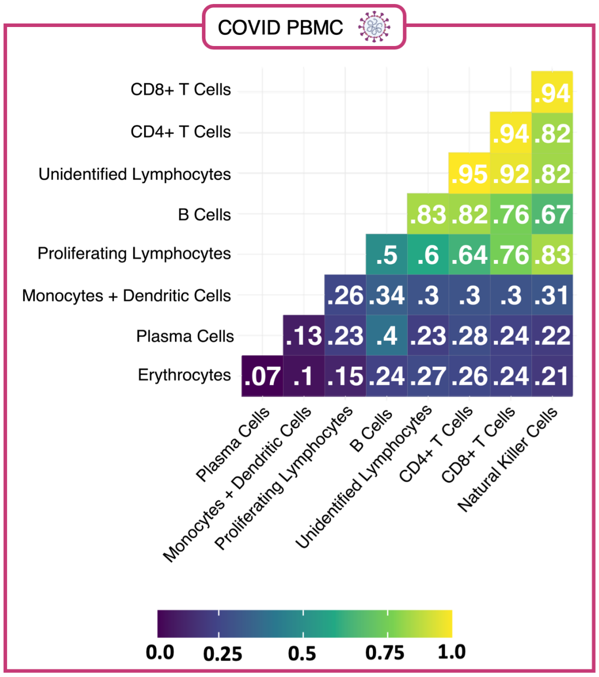


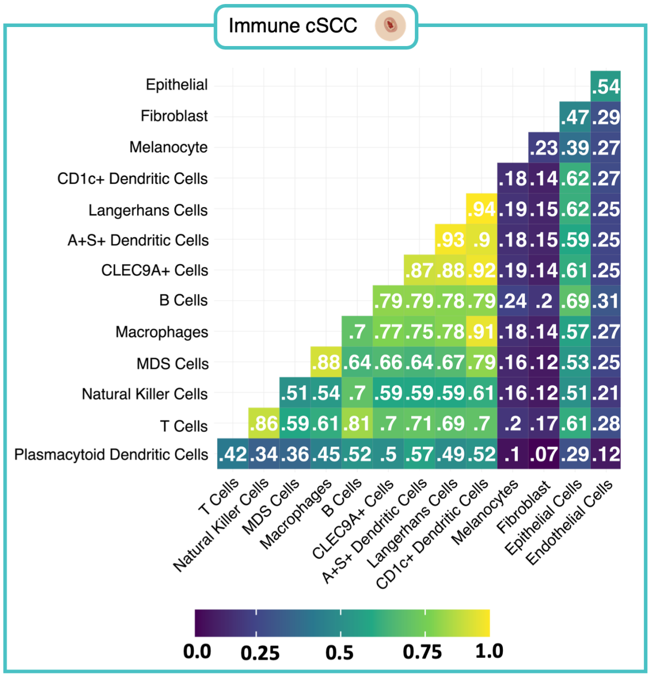


Figure A.5: (For the ease of the reader) An enlarged version of correlation results presented in Fig. 2 of the main manuscript.

### Appendix F. N-ACT Architecture Details

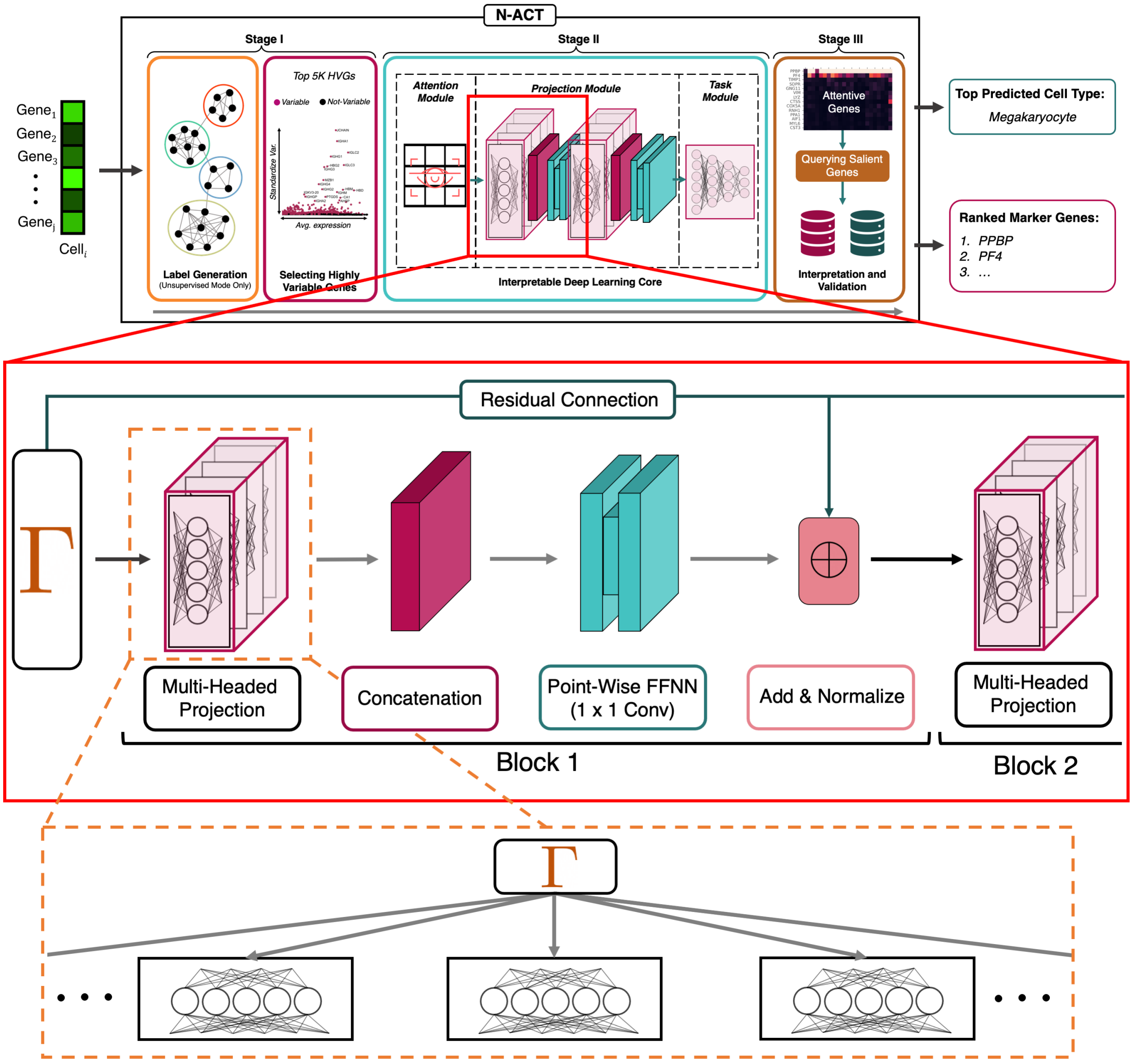


Figure A.6: **A Detailed View of N-ACT and the Multi-Headed Projection Module**. We dive deeper into the projection block of N-ACT in Appendix F. An implementation view of the DL architecture is presented in Fig. A.7.

#### F. 1. Projection Block

The idea behind the N-ACT projection block is to learn various representations for different gene subsets in each cell. Projection block design was inspired by the multi-head attention architecture presented in (Vaswani et al., 2017); however, the projections are not multi-head attention mechanisms. One way to think about the multi-head projection block is to view it as a set of $h$ linear projections, with each $l_h : \mathbb{R}^{B\times N} \to \mathbb{R}^{B\times d}$ ($B$ is the number of samples and $N$ is the number of genes) done sequentially and independent of one-another (*e.g.* in a `for` loop). However, as noted by (Vaswani et al., 2017), these projections can be done more efficiently through creating a tensor $L \in \mathbb{R}^{B\times h\times d}$, which acts the same way as the collection of the individual linear operators. In this formulation, $0 \equiv N (\text{mod } h)$, and the last projection component, the

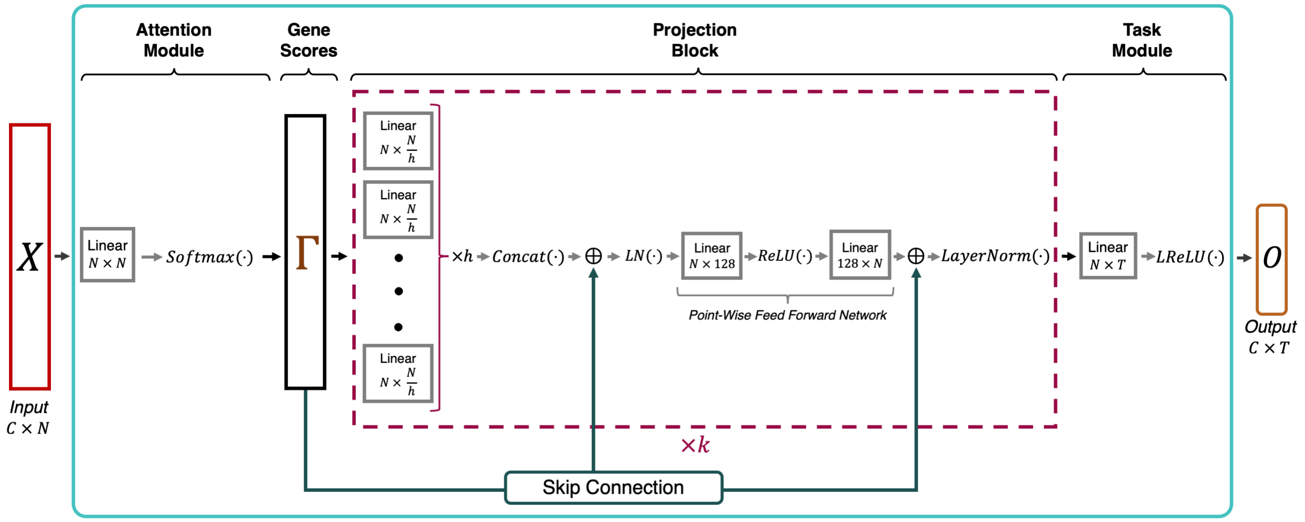


Figure A.7: **An Implementation View of the N-ACT DL Core**. In this illustration, $C$ corresponds to the number of cells inputted, $N$ denotes the number of features ($N = 5000$ in our results), $h$ is the number of projection heads, $k$ is the number of projection blocks(in our model, $h = 10$ and $k = 2$) and $T$ denotes the number of classes (distinct labels) present in the data.

"concatenation" layer (depicted in Fig. A.6) allows us to reshape the model output which can be passed along to remaining layers.

To increase model capacity and allow N-ACT to learn complex mappings, outputs are non-linearly "activated" through a Point-Wise Feed Forward Neural Network (which can be thought of as $1\times$ 1 convolution). However, there is the possibility of projection blocks learning a non-linear mapping unrelated to interpretability, resulting in loss of interpretability. Therefore, we hypothesized that adding residual connection between output of the attention module and input of each subsequent layer would improve performance and interpretability. Our ablation studies show that adding a residual connection improves model performance (Table A.3). Skip connections are added to projection block outputs and normalized using Layer-Norm (Ba et al., 2016)) before being used as input for the next layer.

#### F. 2. Multi-Head Ablation Study

We investigated the effect of head number in the multi-head projection block and found that 10 heads provided the best results (Table A.2). We also tested our model accuracy when using one, two and three projection blocks and found that two projection blocks provided the best balance of accuracy and efficiency. To test our hypothesis regarding residual connections, we tested N-ACT (10 heads, 2 projection blocks) with and without skip connections. As shown in Table A.3, model accuracy drops without the residual connections. Residual connections are identity mappings added to the output of each layer, with this quantity normalized and used as inputs for subsequent layers.

Table A.2: **Ablation Study on the Number of Projection Heads.** We fixed all other hyperparameters and studied the effect of head number on accuracy and interpretability. Training times are the average of 5 training runs on an A100 GPU. Accuracy of each model remained the same across different training settings, since all random parameters were initialized with the same random seed.

| Number of Heads | W-F1 | NW-F1 | Hit@5 | Avg. Training Time |
|---|---|---|---|---|
| 1 | 0.9278 | 0.8968 | 0.85 | $9.11 \pm 0.14$ (Min) |
| 5 | 0.9317 | 0.9077 | 0.85 | $9.38 \pm 0.31$ (Min) |
| 8 | 0.9307 | 0.9156 | **1.00** | $9.31 \pm 0.22$ (Min) |
| 10 | 0.9322 | **0.9173** | **1.00** | $9.56 \pm 0.27$ (Min) |
| 20 | **0.9324** | 0.9156 | **1.00** | $9.87 \pm 0.24$ (Min) |

Table A.3: **Effectiveness of Residual Connections on N-ACT.**

| Residual Connection? | W-F1 | NW-F1 |
|---|---|---|
| Yes | **0.9322** | **0.9173** |
| No | 0.9243 | 0.8725 |

#### F. 3. Training Scheme

As mentioned in the main manuscript, we train N-ACT for 50 epochs using Adam (Kingma & Ba, 2014) optimizer, with a fixed learning rate. We tested training N-ACT for more epochs and found that the accuracy of predictions increases; however, a learning rate scheduling is beneficial in avoiding overfitting (when training over 100 epochs). When training for longer epochs, we employed an exponential learning rate decay with $\gamma = 0.95$ and a decay schedule of every 10 epochs. We chose 50 epochs to balance accuracy and training time efficiency.

#### F. 4. Supervised ACTI Comparison

Table A.4: **Benchmarking N-ACT on Supervised ACTI**. Although the main goal of N-ACT is interpretable ACTI in an unsupervised manner, our model can be used in a supervised setting as well, while still providing biological interpretability. The reported training times are the median of 5 runs for each model. The same random seed was used to initialize all random parameters to ensure reproducibility of results across different runs. *Training/Testing Support*: the number of samples used in training/evaluation, *W-F1*: Weighted F1 score, *NW-F1*: Non-Weighted F1 score

| Model | W-F1 | NW-F1 | Training Support | Testing Support | Med. Training Time |
|---|---|---|---|---|---|
| | | | **Mouse HDF** | | |
| ACTINN | **0.9703** | 0.9677 | 21,003 | 2,998 | **1.5 Minutes** |
| N-ACT (Ours) | 0.9681 | **0.9712** | 21,003 | 2,998 | 5.4 Minutes |
| | | | **Immune CSF** | | |
| ACTINN | **0.9357** | 0.8898 | 66,549 | 8,991 | **3.8 Minutes** |
| N-ACT (Ours) | 0.9285 | **0.8963** | 66,549 | 8,991 | 11.2 Minutes |
| | | | **COVID PBMC** | | |
| ACTINN | **0.9323** | 0.9148 | 55,005 | 9,864 | **3.1 Minutes** |
| N-ACT (Ours) | 0.9322 | **0.9173** | 55,005 | 9,864 | 9.4 Minutes |
| | | | **Immune cSCC** | | |
| ACTINN | 0.9646 | 0.9315 | 40,027 | 6,994 | **2.4 Minutes** |
| N-ACT (Ours) | **0.9684** | **0.9455** | 40,027 | 6,994 | 7.7 Minutes |

### Appendix G. Utility of N-ACT for Disambiguation of Broad Annotations

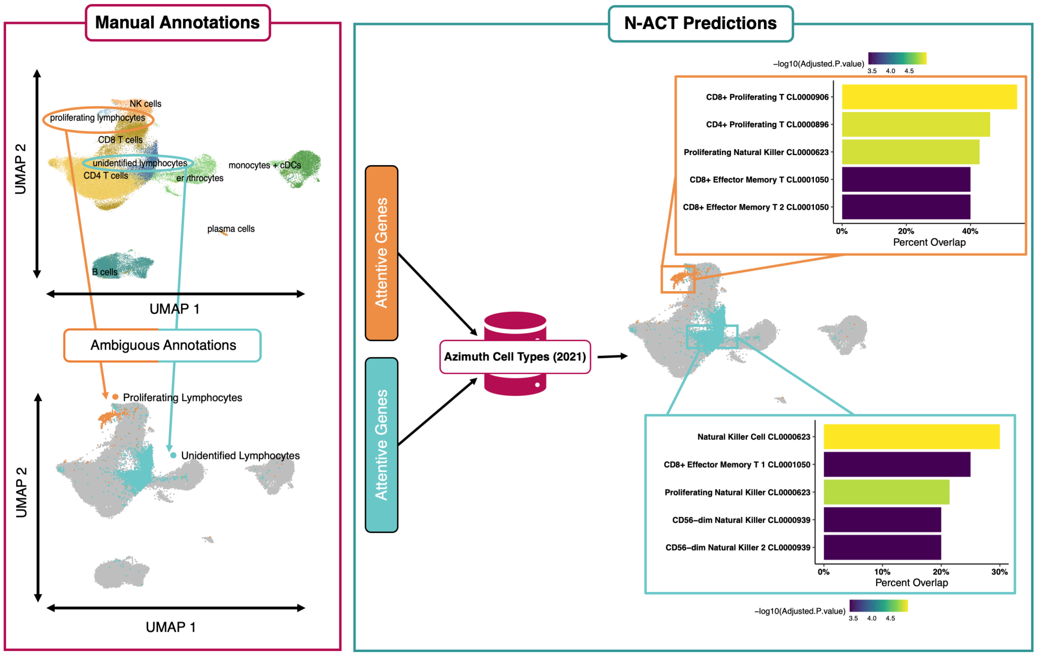


Figure A.8: **N-ACT Used to Disambiguate Broad Annotations**. Here, we use N-ACT-identified salient genes to query a specialized database (Azimuth) to disambiguate original broad manual annotations.

### References (Appendix)


[A1] Ba, J. L., Kiros, J. R., and Hinton, G. E. Layer normalization, 2016. URL https://arxiv.org/abs/1607.06450.

[A2] Butler, A., Hoffman, P., Smibert, P., Papalexi, E., and Satija, R. Integrating single-cell transcriptomic data across different conditions, technologies, and species. *Nature Biotechnology*, 36:411–420, 2018. doi: 10.1038/nbt.4096. URL https://doi.org/10.1038/nbt.4096.

[A3] Chen, E. Y., Tan, C. M., Kou, Y., Duan, Q., Wang, Z., Meirelles, G. V., Clark, N. R., and Ma'ayan, A. Enrichr: interactive and collaborative html5 gene list enrichment analysis tool. *BMC Bioinformatics*, 14:128, Apr 2013. ISSN 1471-2105 (Electronic); 1471-2105 (Linking). doi: 10.1186/1471-2105-14-128.

[A4] Goodwin, S., McPherson, J. D., and McCombie, W. R. Coming of age: ten years of next-generation sequencing technologies. *Nature Reviews Genetics*, 17(6):333–351, 2016. doi: 10.1038/nrg.2016.49. URL https://doi.org/10.1038/nrg.2016.49.

[A5] Hao, Y., Hao, S., Andersen-Nissen, E., Mauck, William M., I., Zheng, S., Butler, A., Lee, M. J., Wilk, A. J., Darby, C., Zager, M., Hoffman, P., Stoeckius, M., Papalexi, E., Mimitou, E. P., Jain, J., Srivastava, A., Stuart, T., Fleming, L. M., Yeung, B., Rogers, A. J., McElrath, J. M., Blish, C. A., Gottardo, R., Smibert, P., and Satija, R. Integrated analysis of multimodal single-cell data. *Cell*, 184(13):3573–3587.e29, 2022/04/30 2021. doi: 10.1016/j.cell.2021.04.048. URL https://doi.org/10.1016/j.cell.2021.04.048.

[A6] Heming, M., Li, X., Räuber, S., Mausberg, A. K., Börsch, A.-L., Hartlehnert, M., Singhal, A., Lu, I.-N., Fleischer, M., Szepanowski, F., Witzke, O., Brenner, T., Dittmer, U., Yosef, N., Kleinschnitz, C., Wiendl, H., Stettner, M., and Meyer Zu Hörste, G. Neurological manifestations of covid-19 feature t cell exhaustion and dedifferentiated monocytes in cerebrospinal fluid. *Immunity*, 54(1):164–175, Jan 2021.

[A7] Heydari, A. A. and Sindi, S. S. Deep learning in spatial transcriptomics: Learning from the next next-generation sequencing. *bioRxiv*, 2022. doi: 10.1101/2022.02.28.482392. URL https://www.biorxiv.org/content/early/2022/03/02/2022.02.28.482392.

[A8] Ji, A. L., Rubin, A. J., Thrane, K., Jiang, S., Reynolds, D. L., Meyers, R. M., Guo, M. G., George, B. M., Mollbrink, A., Bergenstråhle, J., Larsson, L., Bai, Y., Zhu, B., Bhaduri, A., Meyers, J. M., Rovira-Clavé, X., Hollmig, S. T., Aasi, S. Z., Nolan, G. P., Lundeberg, J., and Khavari, P. A. Multimodal analysis of composition and spatial architecture in human squamous cell carcinoma. *Cell*, 182(2):497–514, Jul 2020.

[A9] Kan, H., Zhang, K., Mao, A., Geng, L., Gao, M., Feng, L., You, Q., and Ma, X. Single-cell transcriptome analysis reveals cellular heterogeneity in the ascending aortas of normal and high-fat diet-fed mice. *Exp Mol Med*, 53(9):1379–1389, Sep 2021.

[A10] Kingma, D. P. and Ba, J. Adam: A method for stochastic optimization, 2014. URL https://arxiv.org/abs/1412.6980.

[A11] Korsunsky, I., Millard, N., Fan, J., Slowikowski, K., Zhang, F., Wei, K., Baglaenko, Y., Brenner, M., Loh, P.-r., and Raychaudhuri, S. Fast, sensitive and accurate integration of single-cell data with harmony. *Nature Methods*, 16(12):1289–1296, 2019. doi: 10.1038/s41592-019-0619-0. URL https://doi.org/10.1038/s41592-019-0619-0.

[A12] Kuleshov, M. V., Jones, M. R., Rouillard, A. D., Fernandez, N. F., Duan, Q., Wang, Z., Koplev, S., Jenkins, S. L., Jagodnik, K. M., Lachmann, A., McDermott, M. G., Monteiro, C. D., Gundersen, G. W., and Ma'ayan, A. Enrichr: a comprehensive gene set enrichment analysis web server 2016 update. *Nucleic Acids Res*, 44(W1):W90–7, Jul 2016. ISSN 1362-4962 (Electronic); 0305-1048 (Print); 0305-1048 (Linking). doi: 10.1093/nar/gkw377.

[A13] Luecken, M. D. and Theis, F. J. Current best practices in single-cell rna-seq analysis: a tutorial. *Molecular Systems Biology*, 15(6):e8746, 2019. doi: https://doi.org/10.15252/msb.20188746. URL https://www.embopress.org/doi/abs/10.15252/msb.20188746.

[A14] McInnes, L., Healy, J., and Melville, J. Umap: Uniform manifold approximation and projection for dimension reduction, 2018. URL https://arxiv.org/abs/1802.03426.

[A15] Stuart, T., Butler, A., Hoffman, P., Hafemeister, C., Papalexi, E., Mauck, W. M., Hao, Y., Stoeckius, M., Smibert, P., and Satija, R. Comprehensive integration of single-cell data. *Cell*, 177(7):1888–1902.e21, 2019. doi: https://doi.org/10.1016/j.cell.2019.05.031. URL https://www.sciencedirect.com/science/article/pii/S0092867419305598.

[A16] Vaswani, A., Shazeer, N., Parmar, N., Uszkoreit, J., Jones, L., Gomez, A. N., Kaiser, L. u., and Polosukhin, I. Attention is all you need. In Guyon, I., Luxburg, U. V., Bengio, S., Wallach, H., Fergus, R., Vishwanathan, S., and Garnett, R. (eds.), *Advances in Neural Information Processing Systems*, volume 30. Curran Associates, Inc., 2017. URL https://proceedings.neurips.cc/paper/2017/file/3f5ee243547dee91fbd053c1c4a845aa-Paper.pdf.

[A17] Wolf, F. A., Angerer, P., and Theis, F. J. Scanpy: large-scale single-cell gene expression data analysis. *Genome Biology*, 19(1):15, 2018. doi: 10.1186/s13059-017-1382-0. URL https://doi.org/10.1186/s13059-017-1382-0.

[A18] Xie, Z., Bailey, A., Kuleshov, M. V., Clarke, D. J. B., Evangelista, J. E., Jenkins, S. L., Lachmann, A., Wojciechowicz, M. L., Kropiwnicki, E., Jagodnik, K. M., Jeon, M., and Ma'ayan, A. Gene set knowledge discovery with enrichr. *Current Protocols*, 1(3):e90, 2021. doi: https://doi.org/10.1002/cpz1.90. URL https://currentprotocols.onlinelibrary.wiley.com/doi/abs/10.1002/cpz1.90.

[A19] Yao, C., Bora, S. A., Parimon, T., Zaman, T., Friedman, O. A., Palatinus, J. A., Surapaneni, N. S., Matusov, Y. P., Cerro Chiang, G., Kassar, A. G., Patel, N., Green, C. E. R., Aziz, A. W., Suri, H., Suda, J., Lopez, A. A., Martins, G. A., Stripp, B. R., Gharib, S. A., Goodridge, H. S., and Chen, P. Cell-type-specific immune dysregulation in severely ill covid-19 patients. *Cell Rep*, 34(1):108590, Jan 2021.
